# Feasibility of intravenous injections of pig plasma extracellular particles into rats — an acute study

**DOI:** 10.1101/2024.11.28.625646

**Authors:** Nicolás Cherñavsky, Stepheny C. de Campos Zani, Marcelo A. Mori, Nina T. Zanvettor

## Abstract

**Background:** Extracellular particles (EPs), especially small extracellular vesicles (EVs), extracted from young animals are increasingly being studied in animal models as agents for regeneration and rejuvenation, with studies using EPs from one species injected into another showing no immune reaction. In this study, we aimed to investigate if the injection of Pig Plasma Extracellular Particles (PPEPs) into rats would produce an acute immune or toxic reaction.

**Methods:** Blood from a young pig was collected, PPEPs were isolated by size exclusion chromatography and injected into young male Sprague-Dawley rats, while the control group received a sterile saline injection. After 9 days, the animals were euthanized and their organs were histologically analyzed for signs of cellular damage or immune infiltration.

**Results:** The treated rats showed no signs of acute immunological reaction, behaving normally immediately after the injections and during the 9 days since the first injection. Throughout the trial period, the animals continued gaining weight normally and the histological analysis of their liver, kidney and spleen showed no signs of acute toxicity.

**Conclusions:** PPEPs from young animals do not cause an acute immune or toxic response when injected intravenously into young male Sprague-Dawley rats.

## Introduction

Extracellular particles (EPs) are structures released by the cells into the extracellular space.[1] This umbrella term includes extracellular vesicles (EVs) and non-vesicular EPs.[2] One type of EP that recently attracted considerable scientific interest are small EVs, often called exosomes, which are vesicles measuring up to 200 nm that are involved in intercellular communication.[3] These vesicles carry different cargoes, either inside their membrane, or on their surface, such as RNA, DNA, proteins and lipids.[4] Recently, small EVs from young organisms or cells have been studied due to their potential regenerative and even rejuvenating properties.[4–10] Several sources of small EVs from young origin were tested in these studies, such as the blood of young animals,[11–13] the extracellular medium of cell cultures — usually stem cells — [14–20] or even the amniotic fluid,[21] all of them showing rejuvenating effects in old animals. Importantly, small EVs from old sources showed deleterious effects in young animals.[9]

Recent studies have shown the benefits of small EVs isolated from the blood of young mice in old mice. Sahu et al. showed that EVs from young mice rejuvenated aged cell bioenergetics in vitro and skeletal muscle regeneration in vivo when injected into old mice,[10] Fitz et al. showed that infusions of small EVs from young serum significantly improved age-associated memory deficits in old mice,[11] and Chen et al. showed that the intravenous injection of small EVs from young animals into aged mice extended their lifespan, mitigated senescent phenotypes and ameliorated age-associated functional declines in multiple tissues.[13] Interspecies experiments have also been performed.[12,13,22,23] Chen et al. analyzed the effects of small EVs isolated from the blood of young male human donors when intravenously injected into aged mice and reported an improvement in cognitive deficits and endurance.[13] They also reported that small EVs from young humans substantially rescued the loss of mitochondrial mass and structural integrity in aged hippocampal and muscle cells from mice.[13] In another experiment, Rao et al. studied the effects of EVs extracted from human urine-derived stem cells when injected intravenously into old mice and reported positive effects on spatial learning and memory abilities, enhancement of muscle strength and motor function and reduction of cellular senescence, besides many other positive effects on age-related hallmarks.[23]

In this context, an interspecies study is of particular interest, as Horvath et al. reported a 67% epigenetic rejuvenation — according to the Horvath epigenetic age clock — of old male Sprague-Dawley rats, when injected with a plasma fraction derived from young adult pigs containing small EVs and potentially other EPs. In addition to the reported epigenetic rejuvenation, the old animals presented several health benefits, such as improvement in memory and learning, improvement in biochemical markers (e.g. blood glucose and triglycerides), and reduction in oxidative stress and chronic inflammation. The authors reported that one significant aspect of their experiment was the high concentration of the small EVs-containing preparation injected, which was intended to be about 4 times the concentration of its components found naturally in the blood of young Sprague-Dawley rats.[12,24]

The fact that such a great amount of EPs from pigs was injected into rats and not only caused no immune reaction but brought several benefits warrants further investigation. Thus, in this study, we aimed to investigate the acute effects of the injection of Pig Plasma Extracellular Particles (PPEPs) into rats. Herein, we describe an acute experiment in which PPEPs isolated from the blood of a young pig were injected into young male Sprague-Dawley rats, following the protocols described in Horvath et al.,[12] to evaluate if this would cause an acute immunological or toxic response.

## METHODS

### Blood Collection

The study design is exemplified in **Figure 1**. For the blood collection, we had access to a blood sample from a pig that had been subjected to regular slaughter practice for consumption in the farm Sítio São Luis (Bariri/SP - Brazil). The procedure followed the official Brazilian guidelines to minimize animal suffering. PPEPs were extracted from the blood of the pig — a 7-month-old 80 kg female free-range pig raised on corn and soy animal feed. The pig was a mixed breed between the Brazilian breeds Cuié and Caruncho. The blood was collected after regular slaughter into sterile recipients containing 252 mL of the anticoagulant CPDA-1 solution per 1 L of blood. A total of around 1.25 L of blood was collected. The recipients with blood were stored in a thermos box during transport and kept at around 35° C for 4.5 hours before PPEPs extraction.

**Figure 1.**
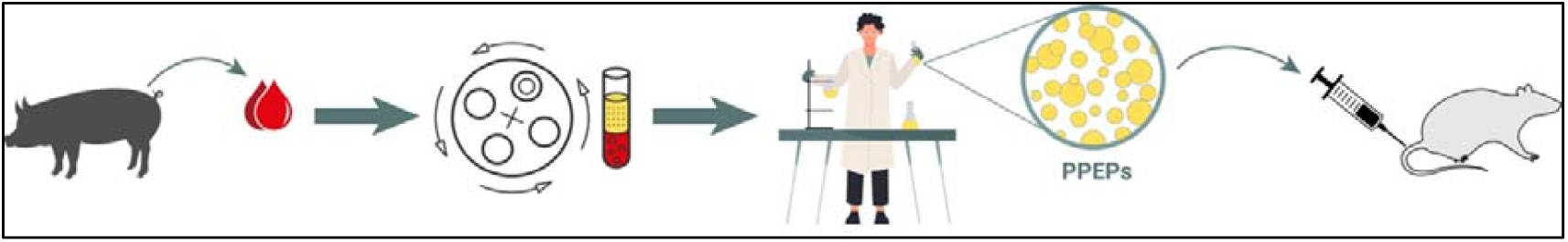
Study design. Refer to text for details.

### PPEPs Extraction

The extraction of PPEPs followed the protocol published by Horvath et al.[12] First, 1 L of the blood was centrifuged at 30° C at 500 x g for 15 min. Then the plasma fraction was carefully collected and pooled using a pipette and further centrifuged at 2,500 x g for 25 min. The resulting platelet-poor plasma was collected carefully using a pipette and added into 50 mL Falcon tubes, with 20 mL of platelet-poor plasma per tube.

To each of these tubes, another 20 mL of a 24% (w/v) PEG 6000 solution in 0.5 M NaCl was added (Sigma Aldrich, St. Louis, Missouri, USA). The combined mixture was incubated for 17 h at 4 °C. After that, the solution was centrifuged at 1,000 x g for 5 min at 4 °C, and the supernatant was discarded. Then, the sticky precipitate was detached from the tube walls using a spatula, and 20 mL of phosphate-buffered saline (PBS) was added into each tube. The tubes were then agitated using Vortex until the precipitate was completely dissolved.

This solution was then filtered in 10 mL size-exclusion columns (Thermofischer, Waltham, Massachusetts, USA) containing Sephadex G-100 (Sigma Aldrich, St. Louis, Missouri, USA) allowed to be hydrated for 48 hours in PBS. The columns were prepared and packed following the manufacturer’s procedure.[25,26] When the columns were ready for use, each column was first washed with 2 mL of PBS, followed by 10 mL of the precipitate solution and then 7 mL of PBS. The first fraction collected (A) was the first 2 mL (EV poor fraction), the second fraction (B) was the following 15 mL (EP rich fraction) and then the third fraction (C) was the last 2 mL (EP poor fraction). Fractions A and C were discarded, and the B fractions were pooled.

The pooled B fractions were then concentrated via overnight dialysis. For this, first, 4 cellulose dialysis tubbing with 12.4 kDa molecular weight cut off (Sigma Aldrich, St. Louis, Missouri, USA) were hydrated in PBS for 1 hour. One end of each tubbing was knotted and tied with cotton yarn. Each tubbing was filled with the pooled B fractions and then the other end was also knotted and tied with cotton yarn. These dialysis bags were then placed in a tray covered with PEG 20000 flakes (Sigma Aldrich, St. Louis, Missouri, USA) and covered with extra PEG 20000 flakes. After 17 hours, the content (PPEPs) inside the dialysis bag was almost dry (semi-solid). The bags were opened, and the content was collected and weighed.

### PPEPs Dosage and Administration

The dosage used for the injections in this study followed the dosage reported by Horvath et al., 572 mg of isolated PPEPs per 200 g rat.[12] The isolated PPEPs were weighed according to the weight of the test animals, and 4 mL of sterile PBS was added to each dose. Then, the tubes were agitated to dilute the PPEPs completely. Each solution was divided into four 1 mL parts, as each animal received the full dose as four 1 mL injections on four different days. The divided doses were stored in a -80 °C freezer.

Six male 5-week-old Sprague-Dawley rats were randomly divided in 2 groups, treated and non-treated (control). The animals in the treated group were injected intravenously into their tail vein with 1 mL of the solution containing the isolated PPEPs, and the other 3 rats received an injection of sterile PBS in the same manner. The four injections happened on alternating days (days 1, 3, 5 and 7), and after each injection, the animals were closely monitored for 20 minutes to assess any sign of acute negative reaction. The rats were weighed on days 1, 5 and 9.

The Sprague-Dawley rats were procured from the CEMIB (Multidisciplinary Centre for Biological Research, UNICAMP, Campinas – Brazil) and were housed during the study in the animal house facility of UNICAMP’s Institute of Biology (Campinas – Brazil) under standard conditions (12:12 h light-dark cycles, 55–70% of relative humidity) at 22 ± 2 °C temperature with free access to water and standard pellet feed (Nuvilab/Quimtia). The experimental protocol was approved by Unicamp’s Animal Ethics Committee. The approval number is 6445-1/2024.

### Tissue Collection and Analysis

On day 9 (2 days after the final injection) all 6 rats were weighed and euthanized by an overdose of isoflurane (> 5%). Their liver, kidney and spleen were collected and a sample was fixed in formalin for further histological analysis. Histological slides were prepared from the paraffin blocks and stained with hematoxylin and eosin. The samples were analyzed via light microscopy for any signs of immune infiltration or structural damage.

## RESULTS

### Effect of interspecies PPEPs injection on body weight

Immediately after each of the 4 injections (on days 1, 3, 5 and 7), all 3 treated rats behaved normally during the 20 minutes of direct observation, and during the 9 days of the experiment. Furthermore, they gained weight at a similar rate to the control rats, as shown in **Figure 2**. There were no signs of distress in any of the groups tested.

**Figure 2.**
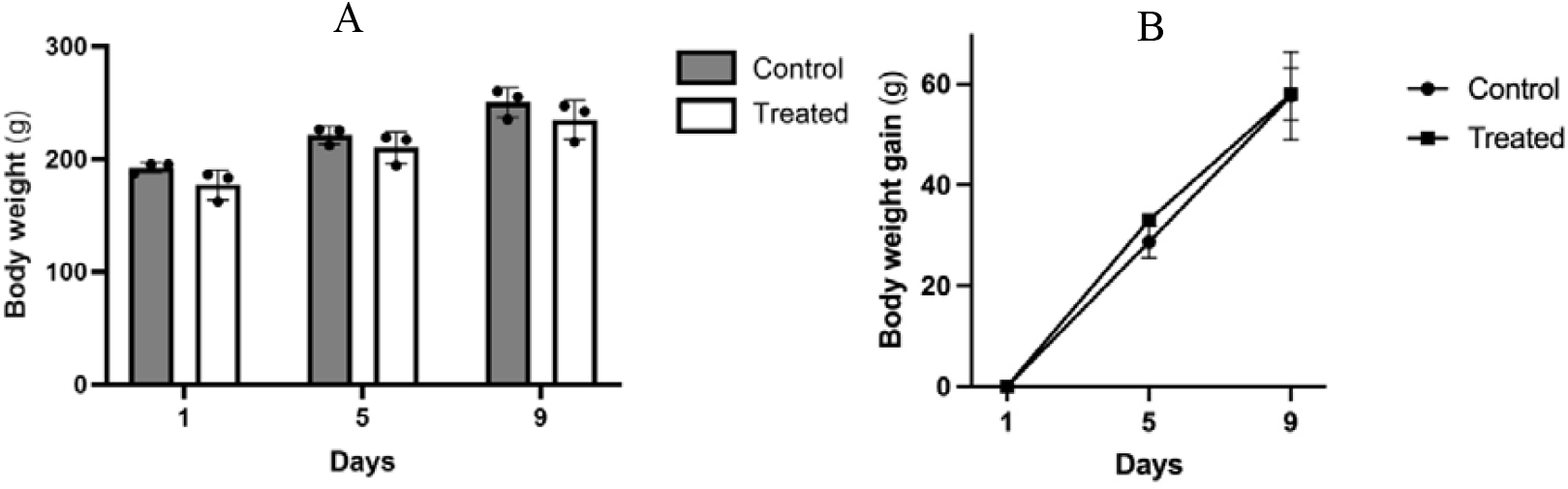
Body weight of treated and non-treated rats throughout the experiment. Data expressed as mean + SD of n = 3 rats per group and analyzed by two-tailed Student’s t-test.Measured body weight (A) and body weight gain compared to day 1 (B).

### Effect of PPEPs on tissue morphology

None of the organs analyzed (liver, spleen and kidneys) showed any signs of acute toxicity or immunological reaction. During tissue collection, the organs showed normal texture, color and size, displaying no signs of edema, infection, or other dysfunction. In addition, there were no changes in tissue architectural structure, immune infiltration or lipid accumulation in any of the tissues analyzed compared to the control group. All tissues presented an integral and well-delimitated structure as healthy tissues as shown in **Figure 3**.

**Figure 3.**
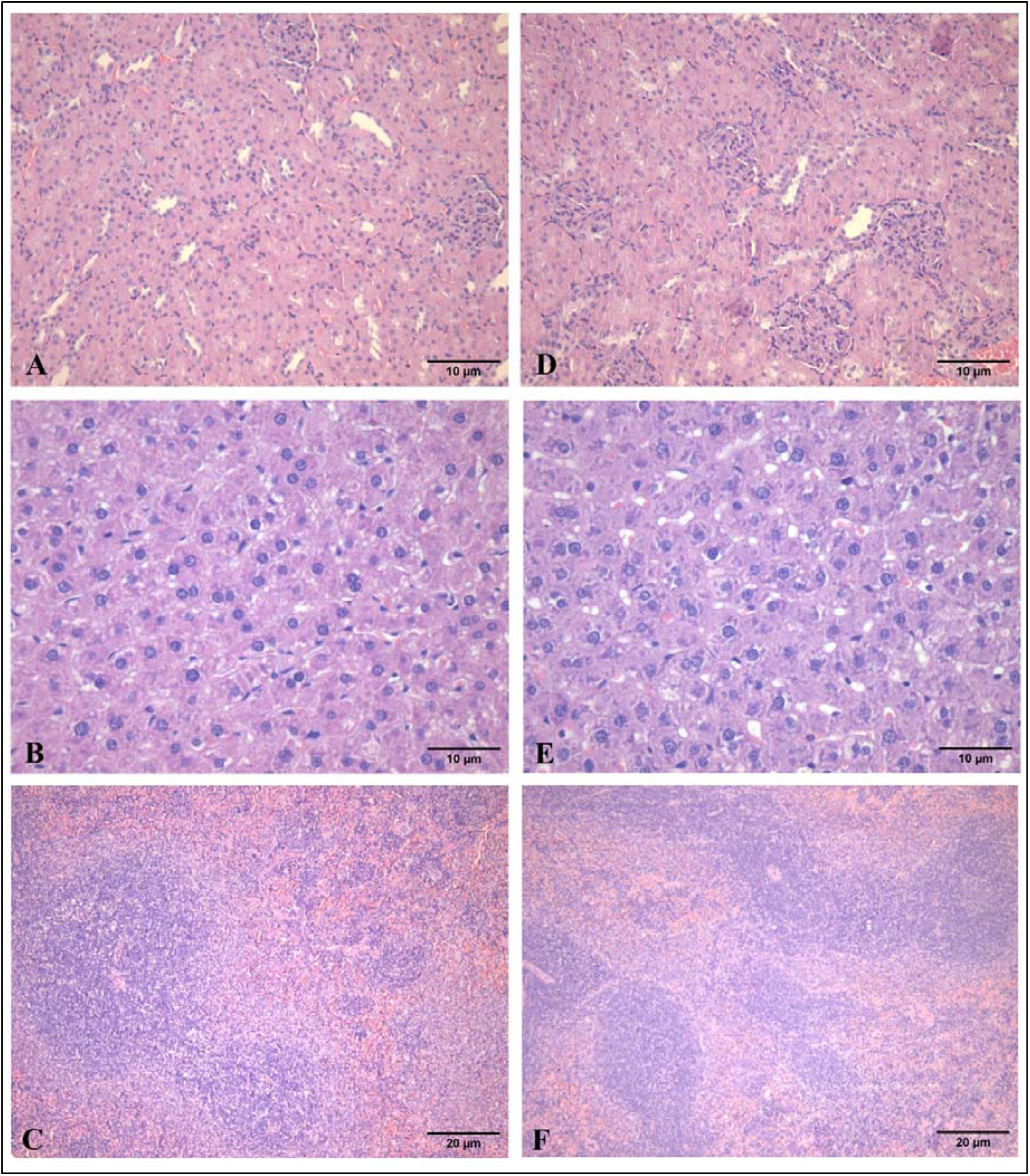
Kidney, spleen and liver of the treated and control rats stained with hematoxylin & eosin. Kidney (A), liver (B) and spleen (C) samples from control rats; kidney (D), liver (E) and spleen (F) samples from treated rats.

## DISCUSSION

The results presented here indicate that the intravenous injection of a high concentration of PPEPs does not induce an acute immune or toxic response in young male Sprague-Dawley rats. Moreover, no changes at the tissue level were observed, suggesting that the interspecies treatment with PPEPs between pigs and rats is well tolerated. However, further research is needed to confirm whether chronic exposure will be also well tolerated.

These results agree with other interspecies experiments described in the literature — one in which small EVs from human blood were injected into mice and another where small EVs from pig’s blood were injected into rats. [12,13] Together, these findings help to strengthen the hypothesis that PPEPs do not induce immune reactions when injected intravenously into a different animal species, at least when both animals are mammals. Nevertheless, more studies with different mammals are necessary to confirm this hypothesis.

Horvath et al. previously showed that it is possible to build a single epigenetic age clock that works for all placental mammal species, indicating that the aging mechanisms at the epigenetic level are very conserved among placental mammals.[27] In fact, several articles published in recent years indicate that signaling factors inside small EVs and in small EVs-containing preparations control and coordinate organismal aging.[12,13,20] The eventual confirmation that small EVs and other EPs do not induce immune reactions in interspecies intravenous injections would be coherent with the idea that a conserved signaling mechanism across mammals regulates aging, and maybe even across other animal classes.[24,28] On that note, Horvath et al. injected a small EVs-containing preparation derived from plasma of young adult pigs into old Sprague-Dawley rats and reported impressive rejuvenation results, such as increased physical strength, increased cognitive function, reduction of oxidative stress and chronic inflammation, and an average of 67% of epigenetic age reduction when compared with the control animals.[12] In the near future, we intend to reproduce their study, as remarkable results demand confirmatory evidence.

One limitation of our study was the number of animals per group (n = 3). This relatively small number of animals was due to the fact that this was a pilot study to investigate the feasibility of injecting PPEPs into rats, before proceeding with a larger study to assess if PPEPs are able to rejuvenate old rats, as reported by Horvath et al.[12], given that, as mentioned, we intend to reproduce these results and further investigate this phenomenon.

## CONCLUSION

We showed that PPEPs do not cause an acute immune or toxic response when injected intravenously in a high concentration into young male Sprague-Dawley rats. However, more studies are necessary to confirm the hypothesis that PPEPs are immunologically safe when used interspecies across mammals and for a prolonged time (chronically). Overall, the experiment described in this article is a first step towards reproducing and further investigating the results reported by Horvath et al.[12]

## ACKNOWLEDGMENTS

We would like to thank Leandro Cardoso, Elzira Elisabeth Saviani and Paulo Augusto de Oliveira Martinez for their technical assistance.

